# The structural basis of cold sensitivity

**DOI:** 10.1101/2025.06.06.658377

**Authors:** Kevin Y. Choi, Xiaoxuan Lin, Yifan Cheng, David Julius

## Abstract

Thermosensitive TRP ion channels enable somatosensory nerve fibers to detect changes in our thermal environment over a wide physiologic range ^1-3^. In mammals, the menthol receptor, TRPM8, is activated by temperatures below ∼26°C and is essential for the perception of cold or chemical cooling agents ^4-6^. A fascinating, yet still unachieved goal is to elucidate structural mechanisms whereby TRPM8 or other thermosensitive channels are gated by changes in ambient temperature. Recent studies using cryogenic electron microscopy (cryo-EM) have attempted to address this challenging question but are limited by difficulties in visualizing temperature-evoked conformational substates or assessing the energetic landscape governing gating transitions ^7,8^. Here, we close this gap by using cryo-EM to visualize TRPM8 channels in cellular membranes, where *bona fide* menthol- and cold-evoked open states are captured. We identify a novel ‘semi-swapped’ architecture in which interdigitation of channel subunits is substantially rearranged following repositioning of the S6 transmembrane helix and elements of the pore region. By combining this structural analysis with thermodynamic measurements using hydrogen-deuterium exchange mass spectrometry (HDX-MS), we are able to pinpoint the pore and TRP helices as key regions undergoing stimulus-evoked conformational dynamics that drive channel gating. Structural mechanisms associated with activation are validated by comparison of human TRPM8 with the menthol-sensitive, but relatively cold-insensitive avian orthologue. We propose a free energy landscape to explain channel gating by cold or cooling agents.

## Introduction

A central goal of structural biology is to obtain ‘snapshots’ of molecules as they transit through conformational and energetic states that define their functional lifecycle ^9^. While advances in cryo-EM have greatly accelerated progress toward this end ^10-12^, there remain numerous challenges and limitations, especially for integral membrane proteins that first need to be extracted from the bilayer using detergents, when auxiliary subunits or other factors required for full biological activity can be lost ^13^. Consequently, some receptor and ion channel structures have only been captured in conformations that likely represent lower energy closed or desensitized states.

This problem is personified by TRPM8, a thermosensory ion channel that is activated by cold, menthol and other cooling agents ^14,15^. Numerous structures of this channel in apo or ligand-bound states have been reported ^16-21^, but thus far none convincingly represent a cold-evoked open state or provide structural insights into thermal gating mechanisms. This problem may be especially pronounced for TRPM8 and other TRP channel subtypes whose biophysical properties suggest the existence of numerous substates associated with multimodal regulation or adaptation ^21-23^. In the case of TRPM8, structural heterogeneity may be exacerbated by the channel’s cold sensitivity, resulting in a range of conformational substates that are too transient or numerous to register as major structural subclasses during cryo-EM analysis (Supplementary Fig. 1). Moreover, difficulties in reconstituting detergent-solubilized TRPM8 protein into vesicles or lipid nanodiscs have limited our ability to observe this channel in its more native membrane environment ^20^. As such, gating mechanisms controlling TRPM8, especially in response to thermal stimuli, remain elusive.

One potential strategy to overcome this roadblock would be to determine TRPM8 structures in cellular membranes, without needing to first extract the channel from the bilayer, as this may provide greater stability for visualizing channels in distinct functional states. Indeed, transmembrane and membrane-associated protein structures have recently been captured in cell-derived vesicles prepared without detergents ^24,25^. By exploiting a similar approach, we identify novel TRPM8 configurations in which large rearrangements within the transmembrane core are associated with stimulus-evoked channel gating, the physiological relevance of which is corroborated by energetic measurements using HDX-MS. Our results reveal the structural basis of TRPM8 cold sensitivity while demonstrating the power of combining structural analysis with thermodynamic measurements to elucidate the conformational landscape underlying physiologic function.

### Novel TRPM8 architecture seen in cell membranes

To visualize TRPM8 in cellular bilayers, we began with an avian orthologue for which there is already detailed structural information ^20,26^. Cells expressing the *Parus major* channel bearing an N-terminal GFP tag were sonicated and inside-out vesicles enriched by affinity chromatography. Our initial analysis was stymied by poor signal-to-noise, which we then mitigated by introducing a second purification step using size exclusion chromatography, yielding micrographs with greatly improved contrast and sample distribution (Supplementary Fig. 2). These raw micrographs showed clear features of TRPM8 in a cellular bilayer which, with further processing, yielded featured 2D class averages that confirmed the presence of cytoplasmic elements facing outward from the vesicle (Supplementary Fig. 2).

From these data, we obtained high resolution (3 – 3.5Å) structures, including the previously described closed and desensitized states ^18,20,27^ (Supplementary Fig. 3), as well as a novel substate whose most notable feature is a distinct domain swap architecture (Fig. 1a). In this configuration, the S6 helix and the associated pore loop are not interdigitated with the neighboring subunit but instead remain associated with their cognate S1-S4 voltage sensor-like domain (VSLD). This novel ‘semi-swapped’ configuration reflects a large repositioning of the S6 helix such that it bends by ∼52° to replace the S6 helix from a neighboring subunit that otherwise engages in the canonical domain swap architecture (henceforth referred to as “fully-swapped”) (Fig. 1b). These changes within the pore region are accompanied by exuberant movements in the associated outer pore loops such that they similarly transition from inter-to intra-subunit interactions, while the relative positions of the pore helices remain unchanged (Fig. 1b). Critical evidence for the semi-swapped configuration is provided by clear densities demonstrating reconfiguration of the connecting regions on both ends of the S6 helix: the pore helix and pore loop of one subunit separate to form the pore domain in conjunction with the neighboring subunit, while the linker between the S6 and TRP helices adopts an acutely bent conformation (Fig. 1c,d).

**Fig. 1.**
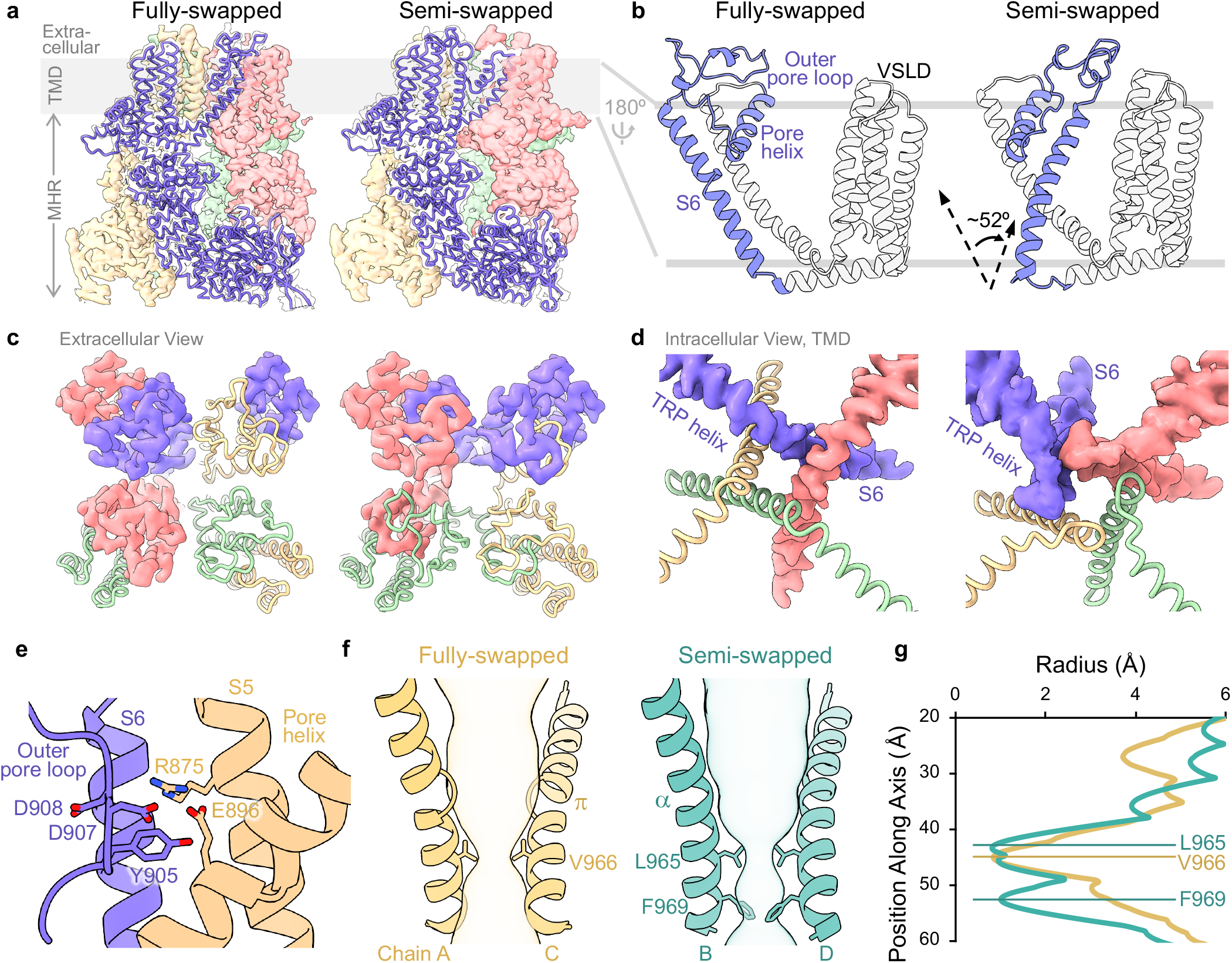
Fully- and semi-swapped configurations of avian TRPM8 in vesicles. (a) Side view of fully-(left) and semi-swapped (right) channel structures determined in cell membrane vesicles. Ribbon diagram of one subunit is fitted into the density. Transmembrane domain (shaded area) and cytoplasmic melastatin homology region are denoted as TMD and MHR, respectively. (b) Reconfiguration from fully- (left) to semi-swapped (right) channel, illustrated for the transmembrane domain of one subunit, is characterized by ∼52° bend of S6 at the junction with the TRP domain, repositioning the outer pore loops, but not the pore helix. (c) Top view showing density and ribbon diagram model of outer pore loop for fully-(left) and semi-swapped (right) configuration. (d) Close up bottom views of S6-TRP helices in fully-(left) and semi-swapped (right) configurations. Subunit colored scheme is the same for panels a, c and d. (e) In semi-swapped configuration, pore domain is stabilized by extensive interactions between Y905 in the pore loop and charged residues in the pore helix and S5 of the adjacent subunit. (f) A closed lower gate constriction is formed by V966 of S6 in the fully-swapped configuration (π-helical form, left panel), or by L965 + F969 in the semi-swapped structure (fully *α*-helical form, right panel). (g) Pore profiles of closed gate in fully- and semi-swapped structure.

Another notable feature of the semi-swapped configuration can be seen at the interprotomer interface, where the S6 helix and outer pore loop of one subunit engages with S5 and pore helices of the neighboring subunit such that Y905 is buried in an amphipathic pocket (Fig. 1e). This novel semi-swapped configuration differs not only from the canonical domain-swapped architecture, but also from previously described non-swapped configurations seen in tetrameric voltage-gated potassium channels ^28,29^. Using the cell-derived vesicle approach, we found that human TRPM8 adopts the same fully- and semi-swapped configurations (Supplementary Fig. 4), demonstrating a conserved architecture across avian and mammalian species.

Notably, in the semi-swapped configuration, a π-to α-helical transition occurs in the vicinity of N958, introducing a register shift by a single residue in the lower half of the S6 helix (Fig. 1f), reminiscent of an agonist-induced gating reconfiguration seen in other TRP channels that is achieved through a p to a transition ^23,30^. Consequently, the lower hydrophobic restriction, formed by V966 in the fully-swapped configuration, is now constituted by L965 and F969 in the semi-swapped state (Fig. 1f,g). This reorientation of the S6 helix is accompanied by repositioning of Y905 at the selectivity filter, thereby widening the upper constriction (Fig. 1g).

### Agonists modulate dynamic equilibrium between states

Among the population of well-resolved particles, we observed a decrease in those corresponding to the fully-swapped configuration (with a concomitant increase in the semi-swapped configuration) in the presence of menthol (Fig. 2a), suggesting that interconversion between these states occurs and is physiologically relevant. To further probe the native state ensemble of TRPM8, we performed HDX-MS experiments to assess both intrinsic and ligand-dependent dynamics of purified, detergent-solubilized avian or human channels. The HDX rate measures the kinetics of exchange between backbone amide hydrogens and solvent deuterium. Exchange is governed by transient breakage of hydrogen bonds during thermal fluctuation, and hence HDX-MS provides a direct readout of protein dynamics and the underlying free energy of unfolding (ΔG) that can be mapped to specific residues ^31^.

**Fig. 2.**
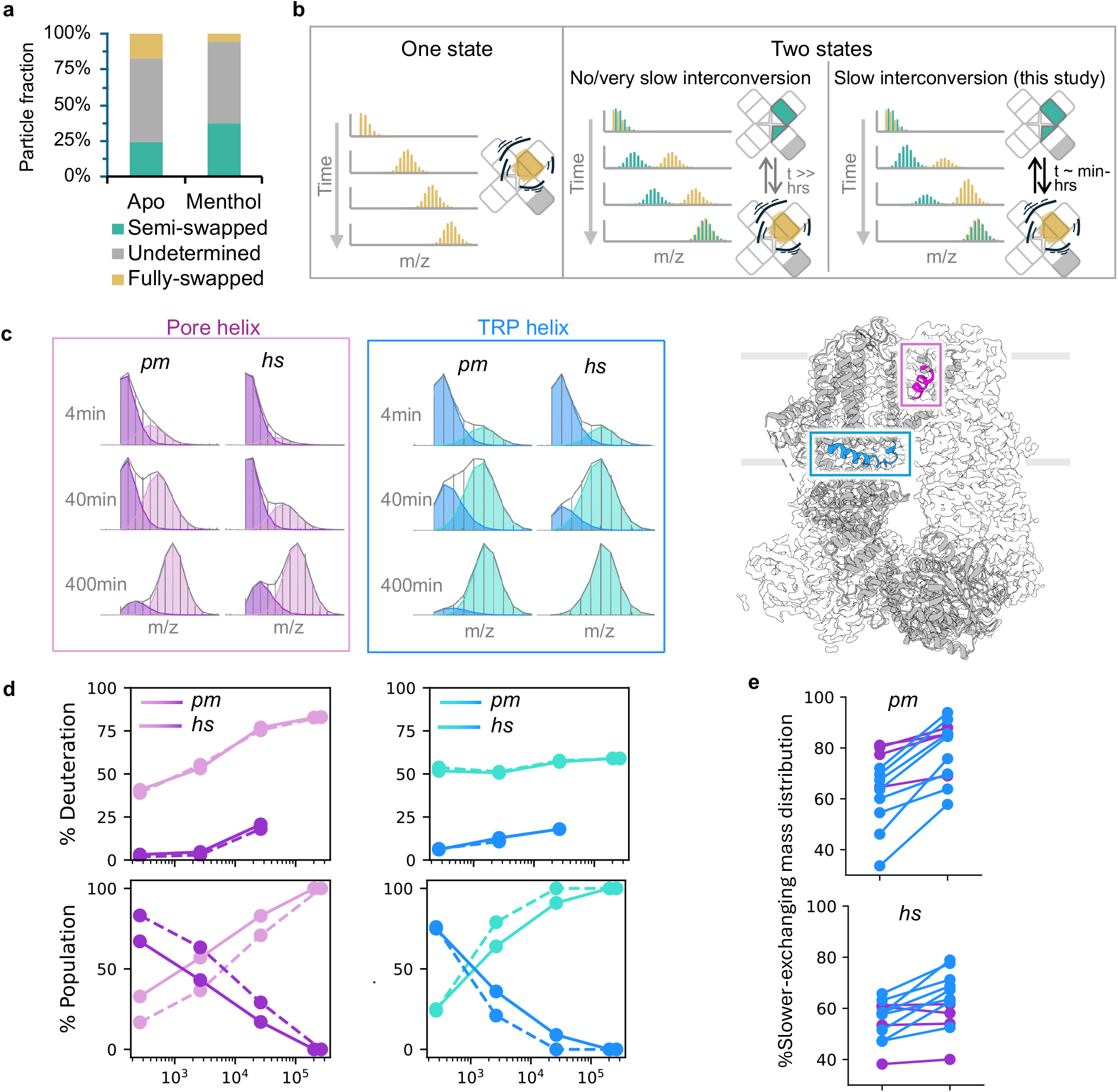
Menthol modulates the dynamic equilibrium of TRPM8. (a) Particle distribution in the apo or menthol-treated pmTRPM8 datasets corresponding to semi-swapped (green), undetermined (grey), and fully-swapped (yellow), configurations. (b) Schematic depicting mass distributions for a deuterated peptide. Left models a “one state” scenario in which peptides with a single conformation or rapid interconversion between conformations show unimodal mass distributions. Right models a “two state” scenario in which peptides with two conformations show bimodal mass distributions, with intensity shifts reflecting interconversion rate between the conformations. (c) Representative mass spectra of peptides from the pore (purple) and TRP (blue) helices (colored in the pmTRPM8 semi-swapped structure, right panel) over HDX times at 4°C for apo avian (pm) and human (hs) channels show bimodal mass distributions (dark/light) characteristic of the coexistence of two slowly interconverting populations. Peptides correspond to residues 891-901 (*pm*) and 901-911 (*hs*) for pore helix, and 982-989 (*pm*) and 992-999 (*hs*) for TRP helix. (d) (Top) Kinetics of deuterium uptake and (bottom) fraction change for the interconverting populations of peptides described in **c**. Solid and dashed lines represent *pm* and *hs*, respectively. (e) Fraction of the slower-exchanging mass distribution (dark purple or dark blue peaks in **c**; presumptive semi-swapped configuration) increases with menthol at 22°C (tHDX=30s), demonstrating agonist-dependent population redistribution.

In these experiments, the majority of peptides exhibited unimodal mass distributions, indicative of a single dominant conformation within the corresponding regions (Fig. 2b). In contrast, peptides covering the pore and TRP helices each showed clear bimodal mass envelopes that increased in mass over labeling time, with one envelope exchanging >100-fold faster than the other (Fig. 2c,d). Such features of bimodality in deuterated mass spectra suggest the coexistence of two conformational populations, with their energetics differing by at least 3 kcal/mol. Notably, the ratio of heavier to lighter mass envelopes progressively increased over labeling time, suggesting that these two populations undergo interconversion on the HDX-MS time scale (i.e., minutes to hours) (Fig. 2b-d).

Importantly, the regions exhibiting bimodality (pore and TRP helices) align closely with those predicted to undergo large conformational transitions between our fully- and semi-swapped models, leading us to propose that the two mass envelopes reflect deuterium uptake associated with these two structural states. Given the additional stabilizing interactions formed at the outer pore region in the semi-swapped configuration of the avian channel (Fig. 1e), we infer that the slower-exchanging population represents the semi-swapped state, while the faster-exchanging population corresponds to the fully-swapped state. This interpretation is further corroborated by the observation that addition of menthol increased the proportion of the slower-exchanging population (i.e. semi-swapped) (Fig. 2e), consistent with the conformational redistribution observed in single particle cryo-EM.

### Agonist stabilizes the TRP helix

We resolved structures of human and avian TRPM8 channels in vesicles in the presence of menthol. These maps enabled us to model the VSLD (the proposed binding site for menthol and other channel modulators) ^19,20,32,33^, thereby revealing a menthol-like density (absent in the unliganded datasets) (Fig. 3a,b). When docked and refined into the cryo-EM density, our models reveal several key interactions between the lower cavity of the VSLD and menthol (Fig. 3b and Supplementary Fig. 5). For example, the hydroxyl group of menthol forms close contact with arginine (R832 in avian or R842 in human TRPM8) (2.9 Å), which is within hydrogen bonding distance. A network of hydrophobic side chains along the inner cavity of the VSLD forms extensive interactions with menthol, likely contributing to its stabilization within the pocket (Fig. 3b). Compared to the closed, unliganded structures, binding of menthol is accompanied by repositioning of the TRP helix such that it is in closer proximity to the VSLD (Supplementary Fig. 5).

**Fig. 3.**
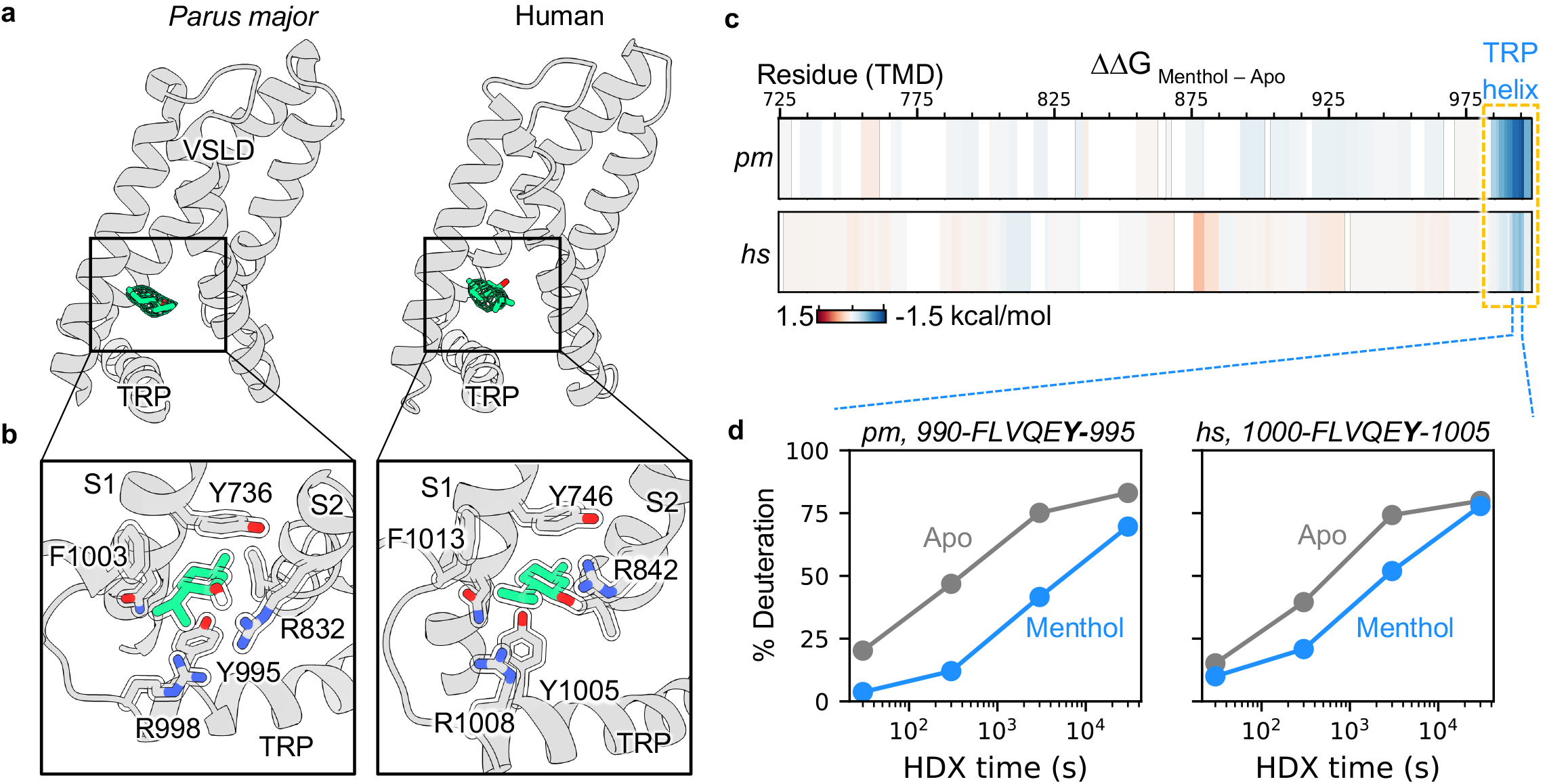
Menthol binding stabilizes TRP helix. (a) Ribbon diagram of VSLD indicating location of bound menthol molecule docked into ligand density (mesh). (b) Atomic details of menthol binding pocket in avian (left) or human (right) channel showing sidechains forming direct interactions with menthol. Residues are annotated. (c) Changes in free energy of unfolding (ΔΔG) for TRPM8 upon menthol binding at 22°C. Residue numbers correspond to the avian sequence. (d) Kinetics of deuterium uptake for the TRP helix indicate that menthol binding stabilizes this region, more so for the avian than human channel.

To investigate the thermodynamic basis driving the menthol-modulated dynamic equilibrium, we performed differential HDX to identify residues exhibiting changes in deuterium labeling in the presence of menthol. Exchange rates for peptides covering the TRP helix slowed by 10-fold, corresponding to a stabilization energy of 1.4 kcal/mol, whereas HDX for the remaining transmembrane domain was unchanged (Fig. 3c,d). This stabilization of the TRP helix includes Y995 (or Y1005 for the human channel), a residue that directly interacts with menthol in our structural models, highlighting the TRP helix as a major energetic contributor to ligand-induced conformational changes. Agonist-induced stabilization was seen whether menthol was added to purified TRPM8 protein or to whole cells prior to protein extraction, indicating that the observed changes reflect intrinsic properties of the channel rather than alterations in cellular or membrane properties (Supplementary Fig. 6).

### Structural basis of differential cold sensitivity

TRPM8 channels show pronounced species-specific differences in thermal sensitivity, with avian orthologues being relatively insensitive to cold compared to their mammalian counterparts (Supplementary Fig. 7) ^34,35^. Despite this functional divergence, the two orthologues share high similarity in both sequence (92%) and structure (RMSD = 1.0 Å for PDB IDs 6O6A and 8BDC). However, our structural analyses in vesicles, as well as HDX-MS measurements, show that the human channel exhibits much greater energetic heterogeneity than previously appreciated. We therefore propose that differential cold sensitivity between mammalian and avian species arises from distinct thermodynamic properties rather than major structural differences in the low-energy states.

Building on this rationale, we sought to identify regions within the mammalian channel that exhibit temperature-dependent dynamics distinct from its avian counterpart. To do so, we performed HDX-MS on avian and human TRPM8 at four temperatures spanning the activation threshold of ∼26°C for the human channel. By converting HDX rates to ΔG’s and folding enthalpy (ΔH) assuming minimal heat capacity changes within each temperature window (37-30°C, 30-22°C, and 22-4°C), we generated sequence-wide, residue-level energetic profiles for both orthologues (Fig. 4a and Supplementary Fig. 8). Although the overall energetics (ΔG) for many regions were similar between orthologues – often differing by less than one k_B_T (spontaneous thermal fluctuation energy) – key differences emerged when examining the enthalpic contribution to folding energy (i.e., ΔH). We found that at 37-30°C and 30-22°C, the human channel exhibited much greater ΔH than the avian channel across the sequence, highlighting the prevalence of temperature-induced dynamics (Fig. 4a). Notably, within the temperature window covering the activation threshold (30-22°C), we observed a markedly slowed HDX at 22°C in two regions in human TRPM8, including the pore helix and the C-terminal helix extending from the TRP domain (Fig. 4b). This HDX slowing and high ΔH (∼ -25 kcal/mol) reflect increased local stability upon cooling, driven primarily by enthalpic contributions. In contrast, cold-induced enthalpic stabilization of these regions was not observed in the avian channel over any temperature range examined, suggesting that they play major roles in conferring cold sensitivity to the human channel.

**Fig. 4.**
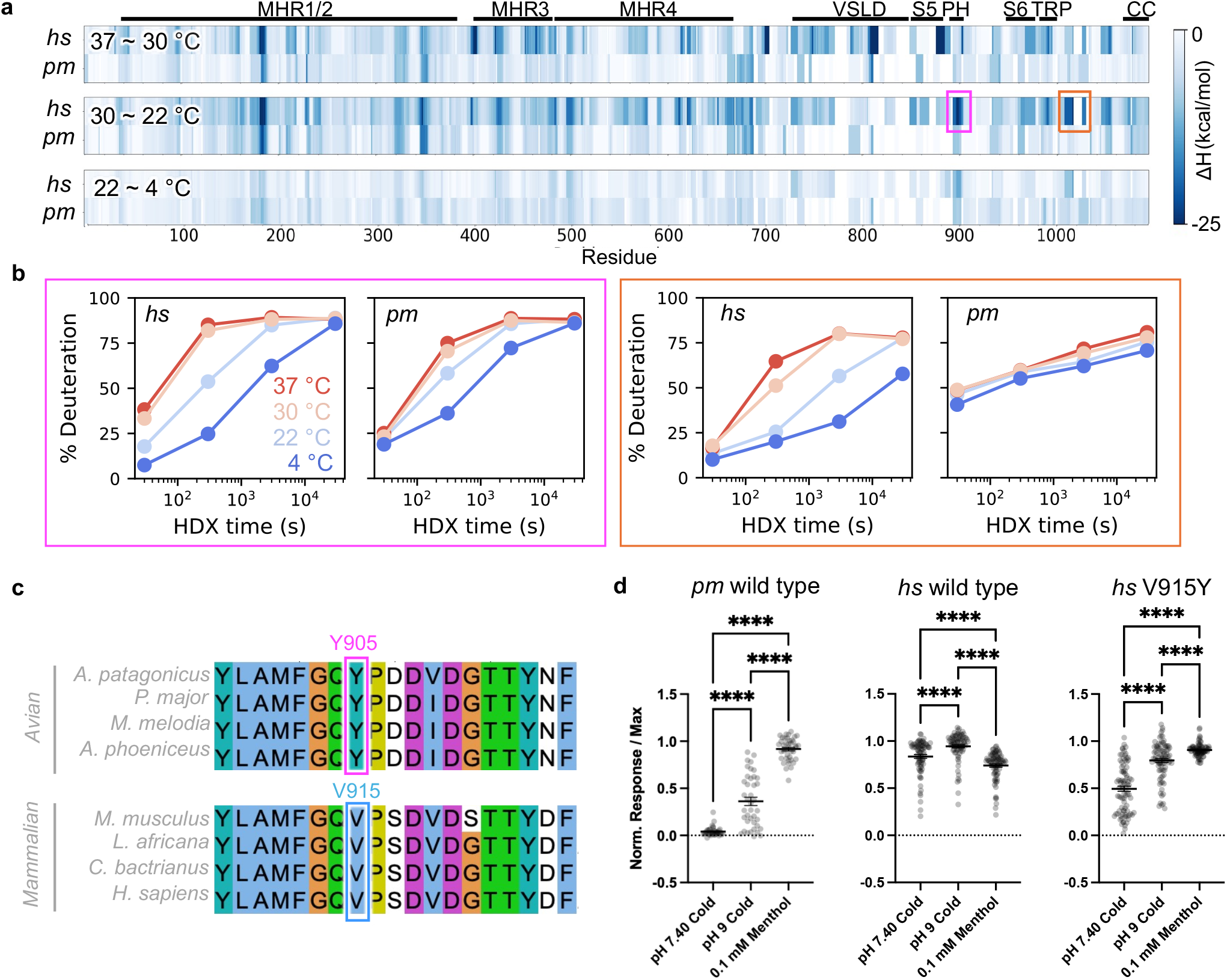
The outer pore region confers species-dependent cold sensitivity. (a) Folding enthalpy (ΔH) for human or avian TRPM8 over three temperature ranges: 37-30°C, 30-22°C, and 22-4°C. Colored boxes for 30-22°C range (spanning activation threshold for the human TRPM8) highlight regions in the human channel with large ΔH that are absent in the avian channel or at other temperature ranges. Residue numbers correspond to the avian sequence. (b) Kinetics of deuterium uptake for representative peptides in regions corresponding to the highlighted boxes in (**a**) at different temperatures. Magenta box: peptide from pore helix, 901-911(*hs*) and 891-901(*pm*); orange box: peptide proximal to the TRP helix, 1016-1026 (*hs*) and 999-1017 (*pm*). (c) Sequence alignment of avian and mammalian TRPM8 at the outer pore region (corresponding to Y905 for the *P. major* channel). (d) HEK293T cells expressing avian or human TRPM8, or the ‘avian-ized’ human V915Y TRPM8 mutant, were exposed to various stimuli, as indicated, and responses measured with Fura-2 ratiometric calcium dye (normalized to maximal calcium response following addition of 10 mM ionomycin). Each dot represents a single cell; n = 40 for pm; 79 for hs and 86 for hs (V915Y). Multi-measure one-way ANOVA with Tukey’s post-hoc analysis; ***P < 0.001, ****P < 0.0001.

The pore helix region is of particular interest given its unique role in forming a stabilizing interface in the semi-swapped (but not fully-swapped) state (Fig. 1e). Interestingly, it has been shown that a single amino acid substitution at this interface, which is conserved as a tyrosine in avian TRPM8 and a valine in mammalian orthologues (Fig. 4c), is linked to species-specific cold sensitivity ^34^. In our structures, we see that this residue is nestled within a pocket lined by a preponderance of charged residues (R875, E896, D907, D908) (Fig. 1e), predicting that this interaction, and thus cold sensitivity of the avian channel, is pH sensitive. Indeed, we found that cold-evoked responses of HEK293 cells expressing the *P. major* channel were markedly enhanced as the pH of the perfusate was increased from 7.4 to 9.0 (Fig. 4d and Supplementary Fig. 7). In contrast, cells expressing the human channel showed robust and consistent cold-evoked responses throughout this same pH range (Fig. 4d). Furthermore, exchanging valine with tyrosine (V915Y) converted cold sensitivity of mammalian TRPM8 to more closely resemble that of the avian channel (Fig. 4d). Indeed, this resulted in a more stable semi-swapped mammalian channel that can be better resolved with cryo-EM (Supplementary Fig. 3).

### A cold-activated TRPM8 structure

Previously described TRPM8 structures report a pore configuration corresponding to presumptive closed or desensitized states, but a clear open state has not yet been described ^16-20^. Our calcium imaging experiments indicate that breaking the species-specific Y905 interface potentiates cold-evoked responses of the avian channel (Fig. 4d). Indeed, under conditions that favor this state (high pH, 4°C), we were able to determine a presumptive open state structure in the semi-swapped configuration using detergent purified channel protein (Fig. 5). Compared to the semi-swapped structure obtained at physiologic pH, where the constriction is formed by two residues (L965 and F969) (Fig. 1f), the S6 helix undergoes an upward translation by one helical turn such that F969 alone now forms the lower constriction (Fig. 5a,b). Concomitantly, the F969 sidechain is reoriented perpendicular to the plane of the membrane, resulting in a widening of the pore from <0.5 to ∼2 Å in radius. Furthermore, we observed a strong cation-like density coordinated in the center of the F969 side chains (Fig. 5a,c and Supplementary Fig. 5). These observations suggest that the above structure determined at high pH and low temperature corresponds to a *bona fide* cold-evoked open state, where F969 can coordinate permeating ions through a π-cation cage-like interaction. Such a lower gate configuration is not unprecedented; for example, the calcium-selective channel TRPV5 forms a permeation pathway involving an analogous tetrameric arrangement of tryptophan residues ^36^. This configuration differs from a previously described open state of mouse TRPM8 obtained in the presence of a cooling compound (cryosim-3) plus an electrophilic agent (allyl isothiocyanate) and soluble di-C8 PIP_2_, in which the gate is formed by a valine residue in S6 with a p-helical configuration ^21^.

**Fig. 5.**
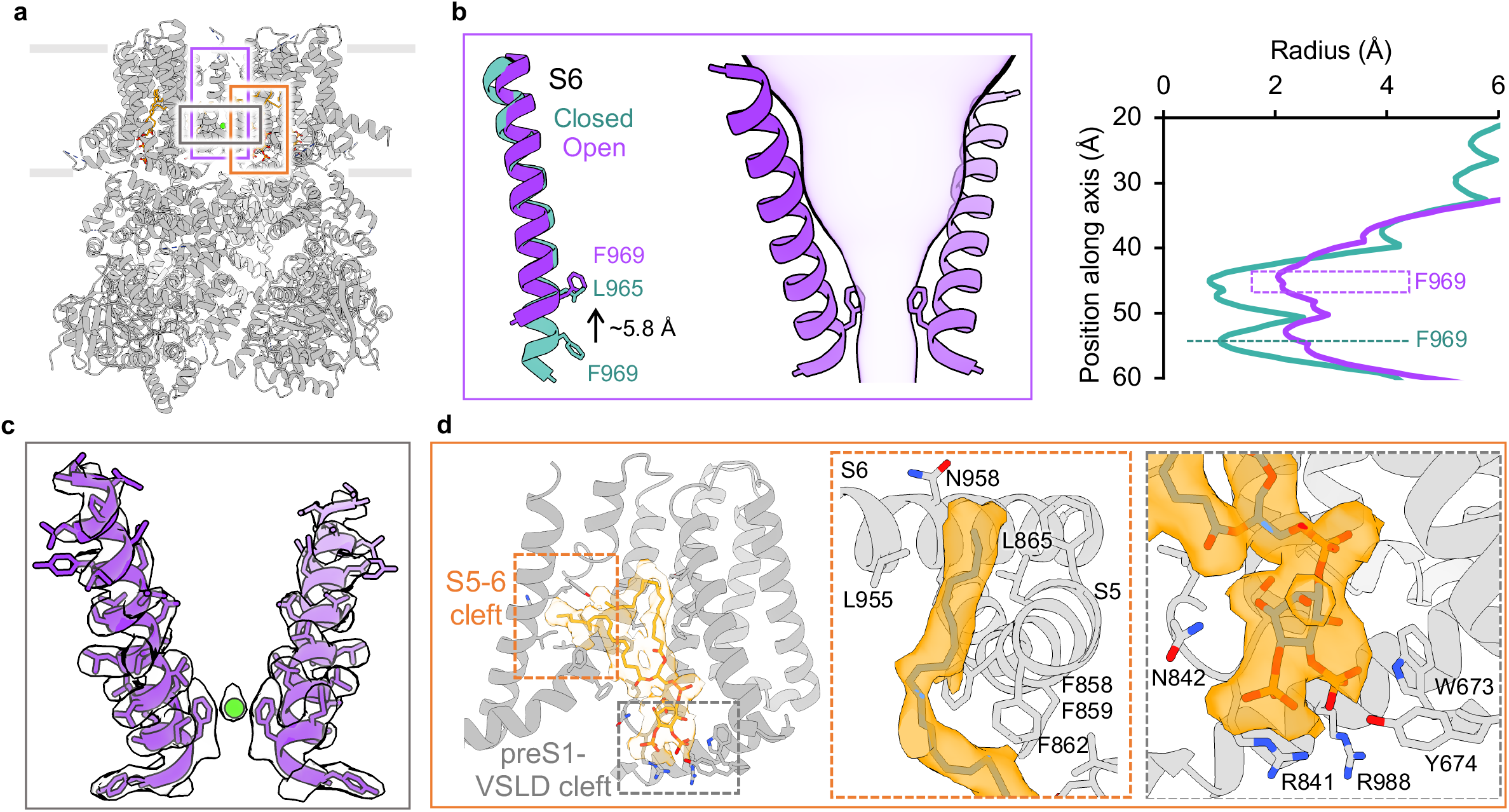
Activation of TRPM8 by cold. (a) Structural overview of pmTRPM8 in semi-swapped configuration with an open pore determined at 4°C, pH 9. (b) The S6 helix in the open semi-swapped structure translates upward by one helical turn when compared to the closed semi-swapped structure; this is accompanied by sidechain rotation of F969 (left). An open lower gate constriction is now formed by F969 (middle and right). (c) Density and model of the S6 helix with a coordinated ion in the open lower gate. (d) PIP2 lipid docked into cryo-EM density with closeups of the hydrophobic S5-S6 cleft and interactions between the PIP2 headgroup and side chains within the pre-S1 and VSLD region.

How is destabilization of the Y905 interface coupled with channel gating? Our structure shows that loss of this interface results in exposure of buried hydrophobic residues at the S5-S6 region that are instead stabilized by a presumptive lipid acyl chain invading this cleft (Fig. 5a,d and Supplementary Fig. 4). The corresponding density connects to a lipid headgroup occupying a previously described binding site for di-C8-PIP_2_ ^19,21^ located at the cleft between the pre-S1 region and the VSLD (Fig 5d). We therefore designate this density as representing a PI(4,5)-bisphosphate lipid, consistent with the finding that endogenous phosphoinositides copurify with TRPM8 in the absence of charged amphiphiles ^21^ and that PIP_2_ is required for channel activation ^37,38^. When modelled, the length of the protruding acyl chain corresponds to 18:0-20:4 PIP_2_, which is the predominant, biologically active species in mammalian plasma membranes ^39,40^. While the position of the S6 helix fluctuates in the absence of stimuli, it is stabilized at a position upshifted by one helical turn by the full-length arachidonyl lipid tail of the endogenous PIP_2_. Consequently, V962 occupies the lipid-contacting cleft in place of N958 and the acyl chain is nestled in a pocket formed exclusively by hydrophobic residues (Fig. 5d). Furthermore, this helical register shift repositions F969 such that the side chain flips upward, disrupting the tight hydrophobic seal seen in the closed state (Fig. 5b,c).

We also determined the structure of human TRPM8 at 4°C in vesicles, where we observed similar features, including an unresolved outer pore loop interface and open gate configuration. In this case, the channel is in the fully-swapped configuration (Supplementary Fig. 4), but S6 adopts the same a-helical conformation seen in the avian open channel as well as a presumptive acyl chain density in the S5-S6 cleft (Supplementary Fig. 4). We hypothesize that because the Y905 interface is absent in the human channel, binding of PIP_2_ occurs more readily at lower temperatures, directly facilitating similar gating rearrangements. At higher temperatures, the intrinsically greater dynamics of the mammalian channel may destabilize lipid binding to prevent opening. Interestingly, the same open configuration was observed for either the avian or human channel in the presence of menthol, suggesting that while cold and menthol act at opposing ends of the S6 helix, the resulting structural rearrangements converge on a common gating mechanism (Supplementary Fig. 4).

## Discussion

Cryo-EM analysis of membrane proteins has evolved to enable the study of receptors and ion channels without having to first extract them from their native environment. In this study, we demonstrate the utility of this approach for capturing conformational states of TRPM8 previously unresolved by us or others using detergent extracted protein. Importantly, we demonstrate that the semi-swapped architecture interconverts with the canonical fully-swapped state in a dynamic equilibrium that is biased by menthol or cold. This was achieved by combining cryo-EM analysis of cell-derived vesicles with energetic measurements of purified proteins using HDX-MS.

Our findings are consistent with an emerging consensus that the dynamic nature of the outer pore and specific protein-lipid interactions are critical factors in specifying the physiological properties of thermosensitive TRP channels ^41,42^. Differential cold sensitivity appears to originate from measurable differences in energetics involving the outer pore loop interface, which is consistent with corresponding structural differences observed for the human and avian channels. This suggests that the outer pore region of mammalian channels evolved with a broader energetic landscape suitable for sensing temperature over a physiologically relevant thermal range. In any case, our structures of avian and human TRPM8 suggest that the functional consequence of outer pore loop dynamics is to regulate accessibility of a hydrophobic tunnel by the long acyl chain of PIP_2_ as a key regulatory factor.

Our results further suggest that opening of the lower gate requires that S6 adopt a full α-helical configuration, which in the avian channel is stabilized in the semi-swapped state. In contrast, a π-helical configuration (seen in previously described desensitized TRPM8 structures) is only observed in the fully-swapped channel structure, likely posing a higher energetic barrier to gating. Thus, we propose that desensitization kinetically traps the channel in this fully-swapped architecture, which may contribute to a process of adaption whereby TRPM8 channels must be returned to temperatures above their thermal activation threshold to achieve full reactivation by cold stimuli ^14^. Furthermore, our structures suggest how PIP_2_ depletion promotes desensitization ^43^ by favoring a return to the fully-swapped state and formation of a p-helical S6 conformation. In any case, a unifying observation is that gate opening requires transition from the p to a configuration, which can occur in the fully-swapped configuration by a single-residue register shift in the lower half of the S6 helix, akin to what is seen in TRPV1, or during transition from fully-to semi-swapped configurations. Together with energetic measurements by HDX, we propose a free energy landscape describing the conformational states observed structurally by cryo-EM (Fig. 6).

**Fig. 6.**
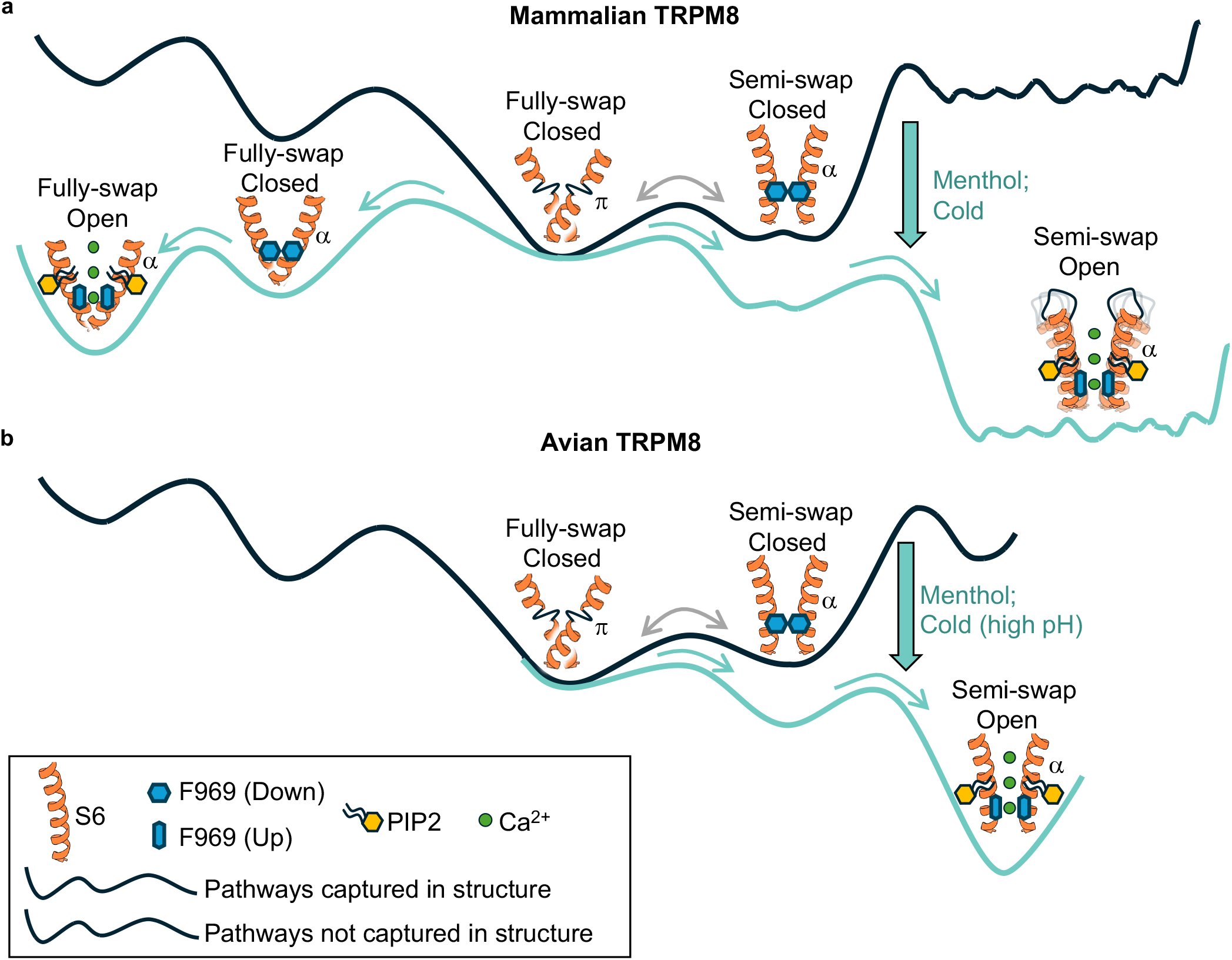
Free energy landscape of TRPM8 activation by cold and menthol. (a) In mammalian TRPM8, the closed channel exists in an equilibrium between fully- (S6 adopts a p helix) and semi-swapped (S6 adopts an a helix) configurations, with the latter occupying a broader energy well compared to the former. Menthol or cold promotes channel opening via two potential pathways: a direct p-to-a transition of the S6 helix in the fully-swapped configuration (left) or transition from fully-to semi-swapped architecture (right). In each case, an a-helical S6 presents a phenylalanine residue at the lower gate, which is stabilized in an upward position to open the pore upon PIP2 binding. The open channel in the semi-swapped configuration exhibits substantial energetic heterogeneity and resides within a particularly broad energy well. As a result, the conformational heterogeneity at the pore region limits its visualization by cryo-EM and only the open structure in the full-swapped configuration is captured.

Our structures suggest that the interconversion between fully-and semi-swapped states requires the movement of pore domains from all four subunits. As such, it is not surprising that reconstructions from a significant proportion of particles in our datasets are characterized by an unresolved pore domain. This phenomenon may explain why previously published ^16^ and numerous unpublished TRPM8 structures from our own studies lack resolution in this region and thus failed to resolve significant conformational changes associated with agonist binding. Such reconstructions are often discarded from further analysis, but our results suggest that they may reflect functionally relevant, dynamic movements. This realization motivates an emerging goal in single-particle cryo-EM, namely, to understand the dynamic processes underlying physiologic function, rather than simply pursuing the highest resolution structures. Furthermore, our findings with TRPM8 suggest that understanding the mechanistic complexity of thermosensation requires a holistic analysis of protein structure and thermodynamics, as we demonstrate here through the combined use of cryo-EM and HDX-MS.

## Materials and Methods

### Cloning, cell culture and protein expression

N-terminally fused mEGFP constructs of TRPM8 were obtained using Gibson assembly to subclone TRPM8 genes into a pFastBac1 vector. Mutagenesis was carried out by either PCR with mutagenic primers, or by Gibson assembly, with Q5 high-fidelity polymerase. Construct sequences were verified using Sanger sequencing and whole plasmid sequencing. Expi293F cells were maintained at 37°C with 8% ambient CO_2_ and grown to a density of 3-3.5×10^6^ cells/ml. To transduce cells for protein overexpression, a baculovirus (generated from two rounds of viral amplification in SF9 cells, according to known protocols) for the N-terminally fused mEGFP *Parus major* or human TRPM8 with a HRV 3C proteolysis site was added dropwise, for a final concentration of 5-10% (vol/vol %) to the flask while gently mixing the culture. Baculoviruses used in mammalian cell cultures were routinely sequenced by amplifying the coding region using PCR (M13 primer sites), gel extracting the amplicon, and subjecting the purified DNA fragments to Sanger sequencing. Virally transduced Expi293F cultures were left at 37°C with 8% CO_2_ for 16 h. To boost expression, the cultures were supplemented with 10 mM sodium butyrate and transferred to an incubator at 30°C with 5% CO_2_ and allowed to incubate for a total of 72 h (post-transduction). Cells were then harvested by centrifugation (3,000 x *g*, 10 min, 4°C), washed once with 1X Dulbecco’s phosphate buffered saline (DPBS) pH 7.40 by gently resuspending the cell pellet, and collected by centrifugation (3,000 x *g*, 10 min, 4°C). Cell pellets were flash frozen in liquid nitrogen and kept in a -80°C freezer until use.

### Detergent purification for HDX-MS and cryo-EM samples

All purification steps were carried out at 4°C or over ice, unless otherwise noted. The avian or human TRPM8 channel was purified with the same general protocol. Briefly, a cell pellet grown from 0.5 L Expi293F cell culture (typically 7-10 g in total) was resuspended into lysis buffer (50 mM HEPES-NaOH pH 7.40, 150 mM NaCl, 50 mg/ml DNase I, 10 mg/ml RNase, 0.2 mM AEBSF, 50 mg/ml soy trypsin inhibitor, 10 mg/ml leupeptin, 10 mg/ml pepstatin, 1 mM benzamidine-HCl, and aprotinin) for a total dilution of 4:1 lysis buffer : cell pellet. The resuspension was adjusted to 0.5% lauryl maltose neopentyl glycol (LMNG) / 0.5% glycodiosgenin (GDN), and gently rotated for 1 h at 4°C. After centrifugation (35,000 x *g*, 30 min, 4°C), the lysate was applied to 1 ml anti-GFP nanobody conjugated Sepharose 4B resin (prepared in-house) for 2 h at 4°C with gentle rotation. The TRPM8-bound resin was washed extensively with wash buffer (20 mM HEPES-NaOH pH 7.40, 150 mM NaCl, 0.05% GDN), then digested in the presence of 10 mg/ml PreScission protease (prepared in house) along with 1 mM DTT for 2 h. The eluate was concentrated to 0.2 ml and injected onto a Superose 6 Increase 10/300 GL column pre-equilibrated in SEC buffer (20 mM HEPES-NaOH pH 7.40, 150 mM NaCl, 0.005% GDN). For the *P. major* structures obtained at pH 9, cells were resuspended into lysis buffer containing 50 mM TRIS-HCl pH 9.0, 150 mM NaCl, 5 mM CaCl_2_, 50 mg/ml DNase I, 10 mg/ml RNase, 0.2 mM AEBSF, 50 mg/ml soy trypsin inhibitor, 10 mg/ml leupeptin, 10 mg/ml pepstatin, 1 mM benzamidine-HCl, and aprotinin. The resuspended cells were extracted using 0.5% LMNG / 0.5% GDN and further purified with anti-GFP nanobody resin. For cryo-EM, the final sample buffer contained 20 mM TRIS pH 9.00, 150 mM NaCl, 5 mM CaCl_2_, and 0.0025% GDN.

### Cell vesicle purification for cryo-EM

A cell pellet grown from 1.0 L Expi293F cell culture (typically 20-25 g in total) was resuspended into lysis buffer (50 mM HEPES-NaOH pH 7.40, 300 mM KCl) supplemented with 2 mM CaCl_2_, 50 mg/ml DNase I, 10 mg/ml RNase, 0.2 mM AEBSF, 50 mg/ml soy trypsin inhibitor, 10 mg/ml leupeptin, 10 mg/ml pepstatin, 1 mM benzamidine-HCl, and aprotinin. The mixture was roughly homogenized using a Dounce homogenizer over ice, and the resuspended cells were adjusted to a final volume of 200 ml. The cells were transferred to a metal beaker over ice and lysed using a probe-tip sonicator (2 min total, 5 s on / 15 s off, 60% amplitude). The lysate was clarified by centrifugation (12,000 x *g*, 15 min, 4°C), and the supernatant was immediately filtered through a 0.8 mm mixed cellulose acetate filter. The filtrate was then passed through 10 ml bed of ion exchange resin (Q-Sepharose FastFlow, Cytiva) packed in a gravity column and preequilibrated in lysis buffer. The flowthrough was collected and adjusted to 0.5 mM fluorinated fos-choline 8 (FFC-8, Anatrace), for a final volume of 240 ml. Vesicles were batch bound to 5 ml anti-GFP nanobody Sepharose 4B resin (prepared in-house) for 2 h at 4°C with constant rotation. After binding, the resin was collected by centrifugation (500 x *g*, 1 min), and transferred into a gravity column. The affinity resin was extensively washed, and the resuspended beads were adjusted to 10 mg/ml PreScission protease (prepared in house) along with 1 mM DTT. Bound vesicles were eluted by proteolysis for 2 h, and the eluate was collected and concentrated using an Amicon ultracentrifugal filter (100 kDa, regenerated cellulose) to a final volume of 0.5 ml. After brief centrifugation (10,000 x *g*, 2 min, 4°C), the sample was injected onto a Superose 6 Increase 10/300 column (Cytiva) preequilibrated with the same elution buffer. The void volume peak (typically 8.5-9 ml) was pooled and concentrated to 30 ml using a 0.5 ml Amicon ultracentrifugal filter (100 kDa, regenerated cellulose) and kept on ice. Concentrated vesicle samples were used immediately for cryo-EM sample preparation.

### Cryo-EM grid preparation, screening, and data collection

Cryo-EM grids for vesicle samples were prepared using either Quantifoil R1.2/1.3 400 mesh holey carbon grids, Quantifoil R1.2/1.3 300 mesh holy carbon grids coated with ultrathin continuous carbon. Detergent samples were prepared using Quantifoil Au R1.2/1.3 300 mesh holey carbon grids. Grids were glow discharged (15 mA, 30 s) and placed in a Vitrobot Mark IV (FEI Company) set at 4°C with 100% humidity. For vesicle samples, 2.5 ml concentrated sample was directly applied to grids, and after 5 seconds of incubation, grids were blotted for 2-5 s and immediately plunge-frozen in liquid ethane cooled in a dewar of liquid nitrogen. To obtain menthol-bound datasets, vesicles were kept at ambient temperature (21-24°C), and a final concentration of 1 mM menthol was added from a stock of 200 mM menthol dissolved in 100% EtOH. After gently mixing the menthol-treated samples and allowing to incubate at room temperature for 1 min, samples were used immediately to prepare grids (for a total incubation time of 2-5 min). For detergent samples, 2.5 ml of purified TRPM8 concentrated to 14-16 mg/ml was directly applied to grids, and after 10 seconds of incubation, grids were blotted for 5 s before vitrifying. Grids were transferred and stored under liquid nitrogen until screening or data collection. Grids were screened with a Talos Arctica or Glacios 200 kV cryo-TEM (Thermo Fisher Scientific) equipped with a K3 direct detector camera (GATAN), and screening datasets were obtained using SerialEM. Data collection was done at the UCSF Cryo-EM Facility with a Titan Krios cryo-TEM equipped with a K3 camera and Bio Quantum post-column energy filter (counting mode pixel size of 0.8189 Å/pix after 2X Fourier binning), or at the Janelia Cryo-EM Facility on a Krios equipped with a cold field-emission gun, Selectris X energy filter, and Falcon 4i camera (physical pixel size of 0.94). For data collections at UCSF, the zero-loss energy selection slit was set to 10 eV. For vesicle datasets, the target defocus was set at –1.0 to –2.5 µm, and for detergent datasets, the target defocus was set at –0.5 to –2.0 µm.

### Cryo-EM data processing, refinement, and model building

Dose-weighted, motion-corrected micrographs were obtained using MotionCor2 ^44^ and Fourier cropped by a factor of two, or by pre-processing using cryoSPARC ^45^. An initial model of *P. major* TRPM8 in a membrane was generated in cryoSPARC with a screening dataset obtained on a Glacios cryo-TEM (2,420 micrographs) and refined to ∼5Å (440 pixel box size, 0.73 Å/pix). This initial reconstruction was used to generate a reference volume using relion_image_handler (512 pixel box size, 0.8189 Å/pix) that was subsequently used in template picking for further processing of datasets obtained on the Krios microscopes. For the vesicle datasets, particles were identified using a combination of Topaz particle picking ^46^ and template picking in cryoSPARC, followed by multiple rounds of heterogeneous refinement and 2D classification to deplete non-TRPM8 or obvious junk particles. After generating a consensus refinement with cryoSPARC non-uniform refinement, further classification was carried out in either RELION5 ^47^ or cryoSPARC. 3D refinement was done with either a mask capturing only the channel region, or with a spherical mask, to limit bilayer signal. Further 2D classification of 3D refinements without a circular mask revealed that a range of membrane curvatures are included in reconstructions, suggesting that in our datasets, 3D classes result from a distribution, rather than specific, membrane curvatures (Supplementary Fig. 2c). For the unliganded *P. major* semi-swapped structure, an initial atomic model was generated *de novo* using ModelAngelo (RELION 5) ^48^, which provided initial coordinates that defined the connectivity of the outer pore loop, S6, and TRP domain. This model was iteratively refined using Phenix refinement and manual refinement in COOT ^49^. For the GDN-purified TRPM8 datasets, an initial reference map was generated from each dataset *ab initio* from particles identified with blob picker in cryoSPARC from a small subset of the micrographs. The *ab initio* model was then subjected to non-uniform refinement without symmetry and subsequently used to generate templates to process the remaining dataset. After multiple rounds of heterogeneous refinement, a consensus refinement was generated in cryoSPARC for further classification with either cryoSPARC or RELION5. Final 3D class averages were refined using either cryoSPARC or RELION5. All models were validated using MolProbity in Phenix ^50^.

### Hydrogen-deuterium exchange and LC-MS

HDX was initiated by diluting 2 μL of 10-25 μM GDN-solubilized TRPM8 stock in an H_2_O buffer into 28 μL of a matching deuterated buffer to reach 93% D-content. Labeling was conducted at pD_read_ 7.0 and temperatures of 4°C, 22°C, 30°C, or 37°C. Labeling times were adjusted to account for the temperature dependence of intrinsic rates (*k*_*chem*_) based on the average *k*_*chem*_ of the full sequence ^51^. For HDX in the presence of menthol, two conditions were employed: 1) 1 mM menthol was added to HEK293 cells expressing TRPM8 and maintained throughout purification and exchange; 2) TRPM8 was purified without menthol, and 1 mM menthol was added only to the D_2_O buffer. Under both conditions, HDX was performed at 22°C.

HDX was quenched at time points ranging from 3 sec to 4 days by adding 30 μL of ice-cold quench buffer (600 mM Glycine, 8 M urea, 0.005% GDN, pH 2.5). Quenched samples were incubated on ice for 20 s, further diluted with 50 μL quench buffer without urea, and immediately injected into a valve system maintained at 5°C (Trajan LEAP). Non-deuterated controls and MS/MS runs for peptide assignment followed the same protocol, except D_2_O buffers were replaced by H_2_O buffers, with immediate quenching and injection. HDX reactions were performed in random order. No peptide carryover was observed as accessed by injecting quench buffer containing 2 M urea and 0.005% GDN. In-exchange controls accounting for forward deuteration towards 41.5% D in the quenched reaction were performed by mixing D_2_O buffer and ice-cold quench buffer prior to protein addition. Maximally labeled controls (“All D”) accounting for back-exchange were performed by incubating samples in D_2_O buffer for 24 h at 22°C, followed by a 1-h heat shock at 48°C.

Upon injection, the protein was digested online using a NepII/pepsin protease column (Affipro AP-PC-006) at 10°C. The resulting peptides were desalted by flowing across a hand-packed trap column (Thermo Scientific POROS R2 resin 1112906 with IDEX C-128 1 mm ID × 2 cm cartridge) at 5°C. Digestion and desalting was completed in 2.5 min at 100 μL/min of 0.1% formic acid. Peptides were then separated on a C18 analytical column (Waters ACQUITY UPLC BEH C18, 130Å, 1.7 µm, 1 mm X 50 mm,186002344) via a 17-min linear gradient of 5–45% (vol/vol) acetonitrile (0.1% formic acid) delivered by a Thermo UltiMate-3000 pump. Eluted peptides were analyzed on a Thermo Q Exactive mass spectrometer in positive mode (full MS: resolution 140,000, AGC target 3 × 10^6^, ma IT 200 ms, scan range 300-2000 m/z; dd-MS^2^: resolution 17,500, AGC target 2 × 10^5^, max IT 100 ms, loop count 10, isolation window 2.0 m/z, NCE 28, dynamic exclusion 15 sec).

### HDX-MS data analysis

Peptides were identified using SearchGUI (v4.0.25) and Byonic (Protein Metrics) against a search library containing TRPM8, 10 additional proteins previously exposed to LC-MS, and their reserved sequences as decoys. Search parameters included unspecific digestion, precursor and fragment tolerance of 10 ppm, precursor charge 1-8, and peptide length 5-50. HDX data were processed in HDExaminer 3.4 (Trajan), with bimodal fitting accepted when the score exceeded the unimodal score by 0.1 or above. The observed exchange occurred via EX2 kinetics, reporting on the equilibrium instead of opening rates of amide protons. For peptides exhibiting bimodal mass envelopes, each envelope increases in mass over labeling time with exchange rates slower than *k*_*chem*_, suggesting that the two mass envelopes represent two kinetically distinct populations that each exchange via EX2 kinetics (rather than EX1 or carryover). To cross-validate those bimodal fits, mass spectra were exported to HX-Express3 ^52^ for double-binomial fitting, with an F-test to access the increase in fit quality over a single-binomial model. Bimodality was accepted when *p*-value < 0.01. Each isotopic distribution was manually inspected for fit quality. Deuteration levels in the uptake curves were adjusted for 93% D-content but not for back-exchange, and those for ΔG calculations were adjusted for both. Except for Figure 2, for peptides displaying clear bimodality, deuteration levels represent centroids of unimodal fit to represent the overall HDX for two populations.

Changes in free energy of unfolding (ΔΔG’s) were calculated from the area between two deuterium uptake curves ^53^. Residue-level ΔΔG’s were determined from a weighted average of peptide-level ΔΔG’s, excluding the first two N-terminal residues due to rapid back-exchange. We note that, while most peptides exhibited near complete exchange within our labeling times, ΔΔG presented here is considered semi-quantitative, as precise quantification requires a broader and denser sampling of labeling times. Nonetheless, this approach effectively captures relative ΔΔG associated with menthol-binding and temperature changes, and our major conclusions remain independent of this fitting method. For Supplementary Fig. 8, a stretched exponential method was used to estimate ΔG from HDX at 22°C ^53^. In those calculations, peptides with exchange rates beyond the measurement range of this study were assigned fixed values: peptides fully exchanged at the first time point (30 s) were assigned *k*_*chem*_=10 s^-1^, while peptides with no exchange at the final time point (30 h) were assigned *k*_*obs*_=10^−5^ s^-1^.

### Fura-2-AM calcium imaging and data analysis

Calcium imaging experiments were carried using Fura-2-AM as a ratiometric fluorescent indicator (as previously described) ^54^ with temperature monitoring throughout imaging. Briefly, adherent HEK293T cells grown at 37°C with 5% ambient CO_2_ were seeded in a 12-well plate (1 ml volume) to a final density of 30-40% confluence and transfected with 0.5 mg/ml plasmid using Lipofectamine 3000 (Thermo Fisher Scientific) according to the manufacturer. Cells were allowed to transfect for 12-14 h. Borosilicate cover slips (12 mm, Bellco Glass) were incubated with a 1:100 solution of Matrigel matrix (Corning) prepared in serum-free DMEM (Gibco) for 1 h, then washed once with DMEM, and left in a 24-well plate with DMEM supplemented with 10% bovine calf serum. Transfected cells were gently dissociated and replated directly onto coverslips and allowed to attach for 2-3 h before imaging experiments were carried out. Attached cells were washed once with Ringer’s solution (10 mM HEPES-TRIS pH 7.40, 140 mM NaCl, 5 mM KCl, 2 mM CaCl_2_, 2 mM MgCl_2_, 10 mM glucose; 290-310 mmol/kg^-1^), and then incubated for 0.5 h at room temperature in the dark in Ringer’s solution supplemented with 10 mg/ml Fura-2-AM (Thermo Fisher Scientific) and 0.02% Pluronic F127. After incubation, cells were washed once with Ringer’s solution and allowed to further incubate in Ringer’s solution without Fura-2-AM or Pluronic F127 for 0.5 h. Dye-loaded cells were then imaged using an inverted microscope with 340 and 380 nm excitation (Sutter, Lambda LS Illuminator). The temperature for each experiment was held at 32-37°C by perfusing Ringer’s solution through a Peltier device controlled with a heating module (Warner TC-324B) calibrated to continuously hold the desired temperature. Cells were held at this temperature range for 5-10 minutes prior to each recording. Bath exchange into low temperatures was achieved by perfusing Ringer’s solution cooled with ice water into the recording chamber. Temperature was monitored and synchronized using a thermistor reading input to the heating module that was digitized through an Axon Digidata 1550B module and monitored with pCLAMP10. Recordings were obtained as stacked movies of individual 340/380 frames with a 1 s interval in Micro-Manager ^55^. Movies of ratio images were calculated with an ImageJ macro script according to known procedures ^56^. Peak amplitudes of calcium signals corresponding to either cold or menthol, were obtained and normalized to maximum calcium signal obtained using 10 µM ionomycin prior to further statistical analyses.

## Acknowledgements

We thank S. Marqusee (UC Berkeley) for providing access to HDX-MS instruments and Byonic software, D. Bulkley, G. Gilbert, L. Wang (UCSF EM Core), R. Yan, and N. Spellmon (Janelia Cryo-EM Facility) for their advice and assistance with cryo-EM data acquisition, and W. Choi for advice with data interpretation and model building. This work was supported by grants from the NIH (R35NS105038 to D.J. and R35GM140847 to Y.C.). Instruments at the UCSF Cryo-EM facility are partially supported by grants from the NIH (S10OD020054, S10OD021741 and S10OD026881) and Howard Hughes Medical Institute. Y.C. is an Investigator of the Howard Hughes Medical Institute.

## Author contributions

K.Y.C. carried out biochemical, structural, and live-cell imaging components of the study. X.L. carried out HDX-MS components of the project. All authors participated in data analysis and evaluation and manuscript preparation. D.J. and Y.C. provided advice and guidance throughout.

---

**Supplementary Fig. 1.**
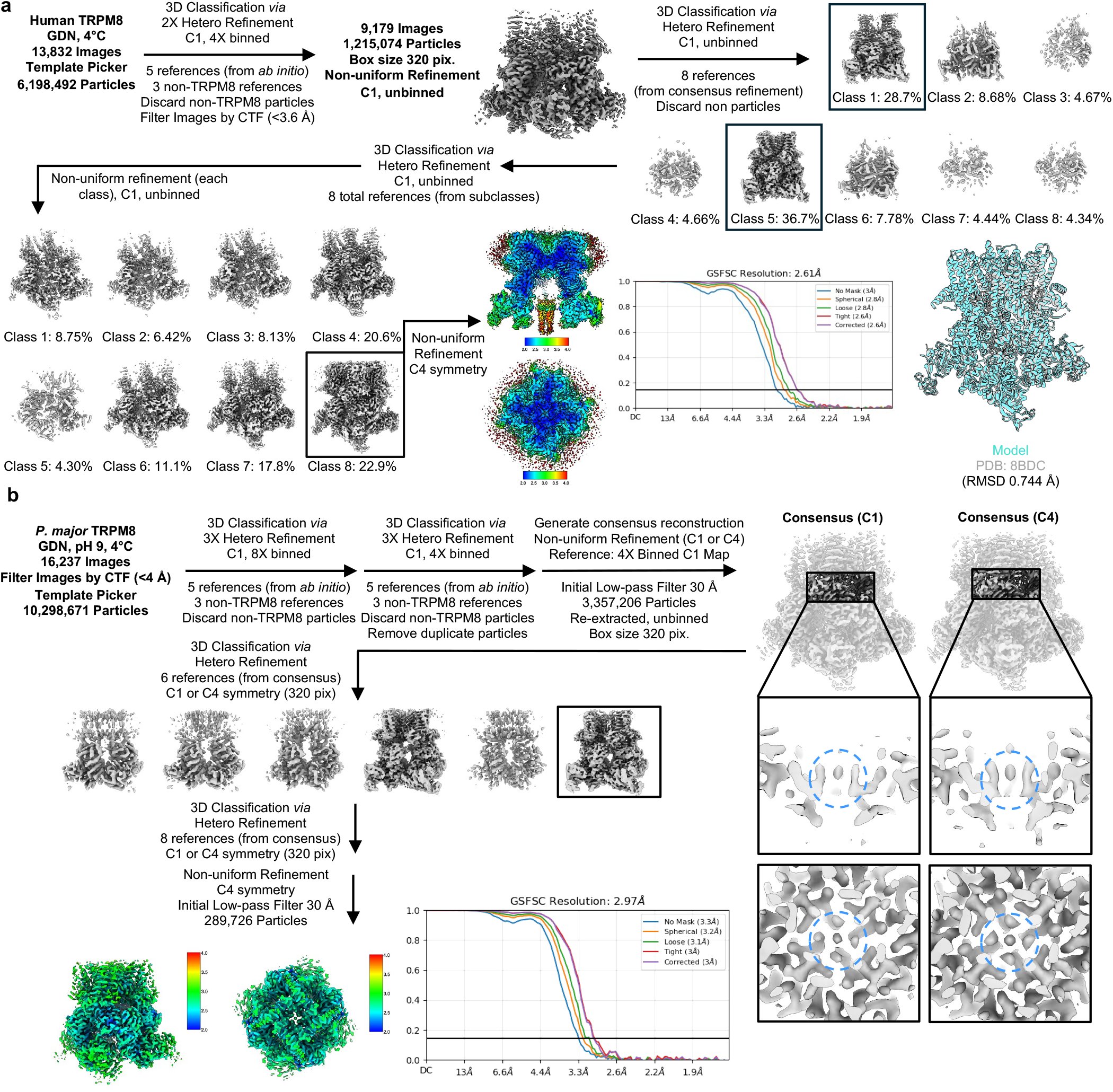
Data processing of TRPM8 structures determined in detergent. (a) Overview of data processing for a GDN-purified human TRPM8 structure determined at 4°C. The consensus refinement is characterized by a well resolved cytosolic domain and poorly resolved transmembrane region. Subsequent classification led to 3D classes with similarly poorly resolved transmembrane regions (bottom right: classes 1-7), and one class with a fully resolved transmembrane domain (bottom right: class 8). Refinement of class 8 with C4 symmetry yielded a map at ∼2.6 Å resolution. An overlay of the fitted model and published hsTRPM8 model (PDB: 8BDC) is shown. This structure resembles the published closed, fully-swapped *pm*TRPM8 structure (PDB: 6O6A; rmsd 1.781 Å), or the closed, fully-swapped *hs*TRPM8 structure (PDB: 8BDC; rmsd 0.744 Å). (b) Overview of data processing for a GDN-purified *P. major* TRPM8 structure determined in high pH at 4°C. Consensus refinement with either C1 or C4 symmetry revealed a lower gate configuration that involves F969 and contains a cation-like density. The displayed local resolution maps, GSFSC plots, and orientation plots were generated using ChimeraX and cryoSPARC.

**Supplementary Fig. 2.**
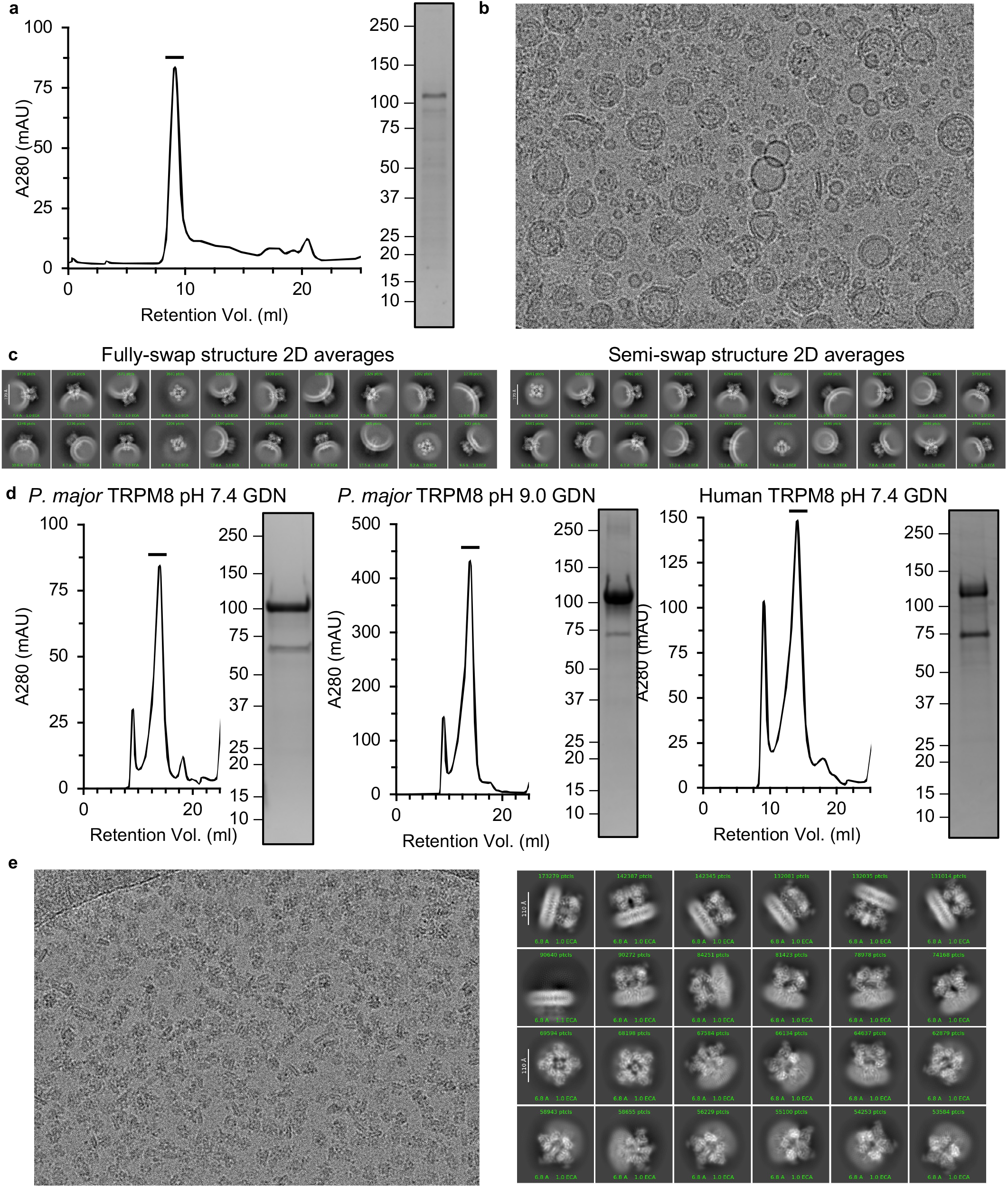
Preparation and cryo-EM of TRPM8. (a) Representative size-exclusion chromatogram of TRPM8 in cell membrane vesicles. Black bar denotes representative region analyzed by Coomassie-stained SDS-PAGE gel. (b) Representative micrograph of TRPM8-containing vesicles. (c) Representative 2D class averages of TRPM8 in membrane vesicles from particles classified as fully- or semi-swapped channels as indicated, in each case showing a wide spread of membrane curvatures. (d) Purification of *pm*TRPM8 or *hs*TRPM8 in GDN. (e) Representative micrograph and 2D class averages of GDN-purified *pm*TRPM8 at high pH.

**Supplementary Fig. 3.**
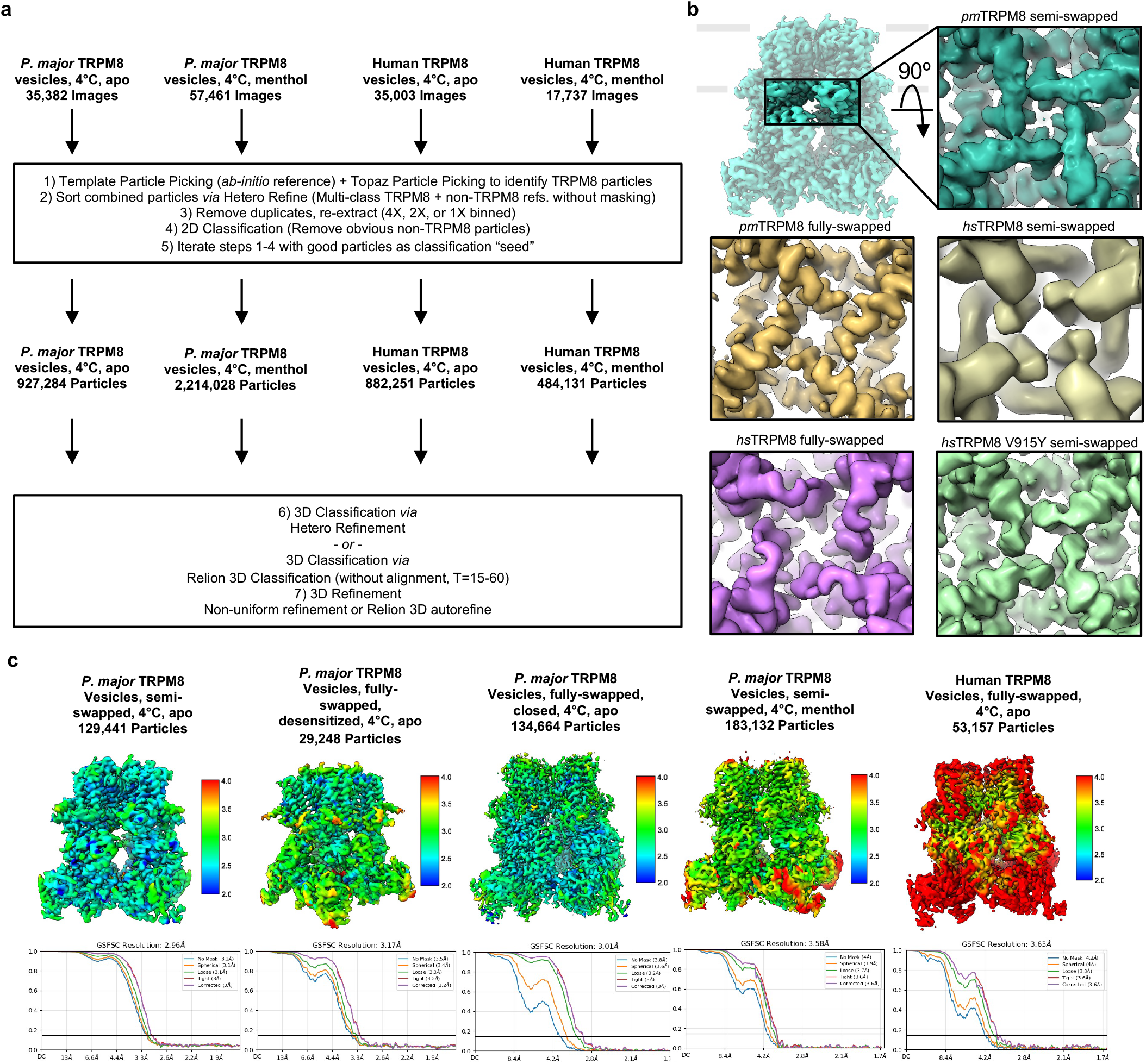
Data processing for TRPM8 in cell vesicles. (a) Overview of data processing pipeline for TRPM8 structures determined in cell vesicles. (b) Bottom view of the fully- and semi-swapped channel seen in avian or human TRPM8 reconstructions. “Avian-ized” human channel (V915Y) shows increased resolution of the semi-swapped configuration, suggesting that the V915Y mutation stabilizes the semi-swapped configuration of the human channel. (c) Local resolution maps and GSFSC curves of TRPM8 structures determined in cell vesicle datasets. Local resolution maps and GSFSC plots were calculated in cryoSPARC.

**Supplementary Fig. 4.**
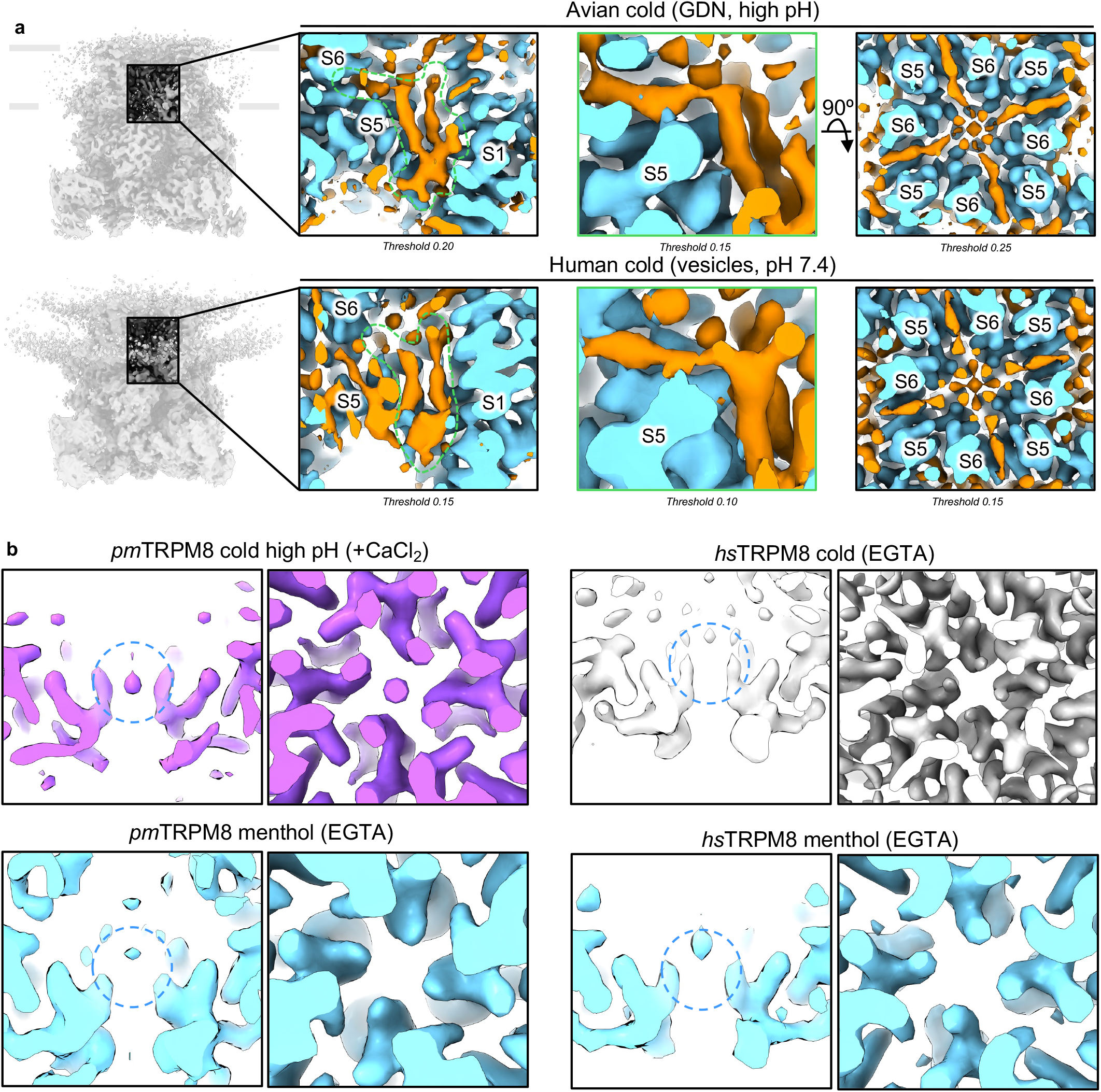
Lipid and ion densities in the cold-activated avian or human TRPM8 structures. (a) Details of lipid density observed at the TRPM8 PIP_2_ binding site for the avian channel determined in cold at high pH or the human channel determined in cold in membranes at pH 7.4. Lipid densities are colored as orange, and modelled protein densities are colored as light blue. (b) Cross-sectional top or side views of the central transmembrane cavity near the lower gate in the presence or absence of calcium.

**Supplementary Fig. 5.**
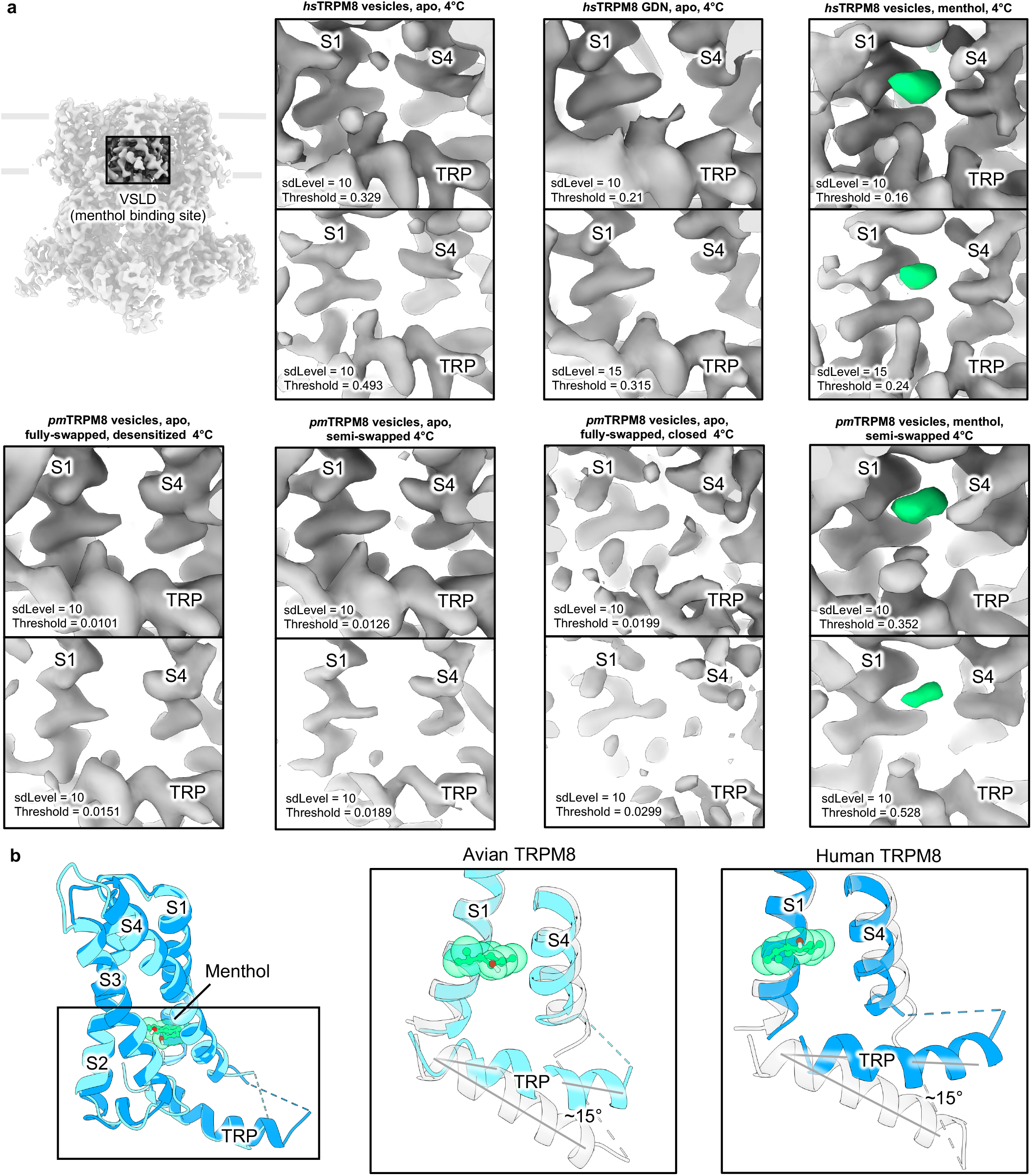
Menthol binding site. (a) Representative structures determined in the presence or absence menthol for the avian or human TRPM8, and close-up view of the VSLD at two map thresholds. Standard deviation levels were set using ChimeraX, and exact map thresholds are displayed. (b) Overview and overlay of the menthol-bound VSLD of avian (dark-blue) or human (light-blue) TRPM8. RMSD was 0.675 Å for 110 pruned atom pairs, or 2.110 Å for all 132 pairs. (c) Relative repositioning of the TRP helix observed between the menthol bound (blue) and unliganded, closed structures (ghosted). In the presence of menthol, the TRP helix repositions by a ∼15° upwards tilt towards the VSLD.

**Supplementary Fig. 6.**
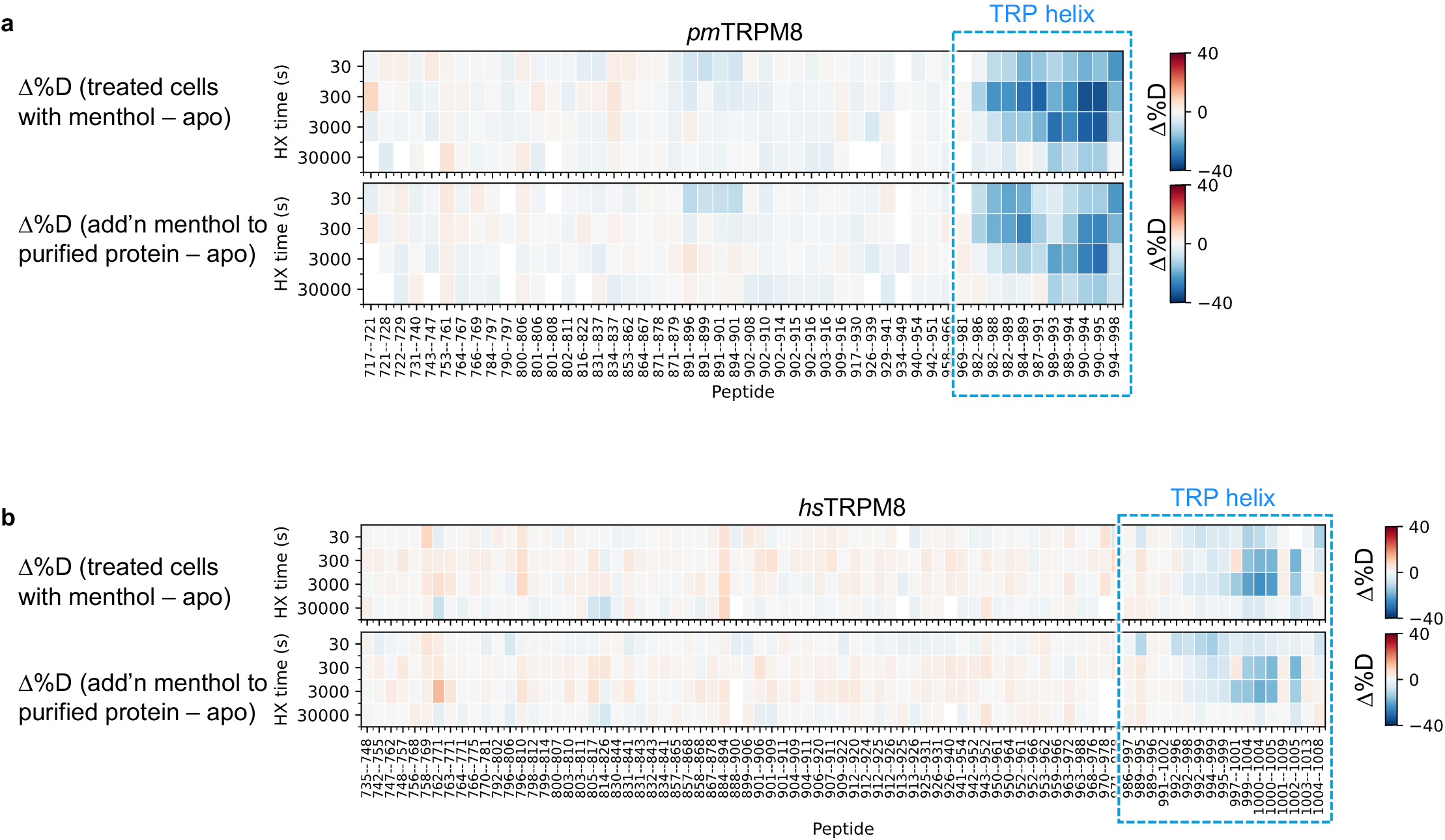
Menthol binding stabilizes the TRP helix. Heatmaps showing changes in deuteration levels at each labeling time for all TMD peptides of avian (a) or human (b) TRPM8 with or without menthol measured in one of the following two conditions: (Top) 1 mM menthol was added to whole cells prior to protein extraction and maintained throughout purification and exchange; (bottom) 1 mM menthol was added to purified TRPM8 protein only during exchange.

**Supplementary Fig. 7.**
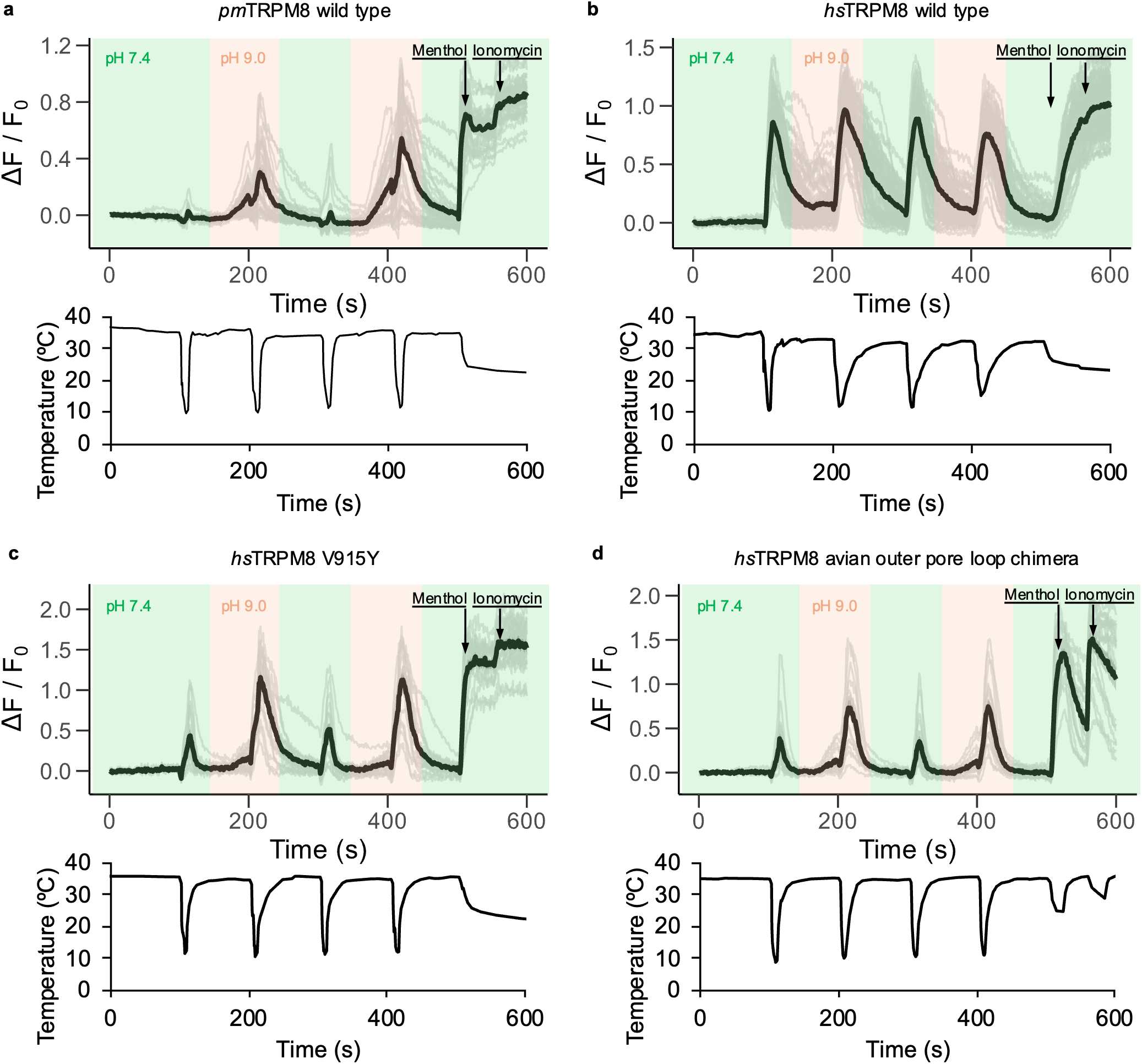
Cold response of avian or human TRPM8. Representative calcium imaging traces obtained from HEK293T cells expressing (a) *pm*TRPM8 wild type, (b) *hs*TRPM8 wild type, (c) *hs*TRPM8 V915Y, and (d) a *hs*TRPM8 chimera replacing the entire outer pore loop sequence of *pm*TRPM8. Bolded traces represent the average response obtained from all measured cells, and light traces represent responses of individual cells. Calcium responses were measured with cell-permeant, ratiometric dye Fura-2-AM. Temperature ramps are shown below each trace. Responses to cold or menthol (100 µM) were normalized to maximum calcium signal following application of 10 µM ionomycin. For figure 4, the first two cold responses were quantified.

**Supplementary Fig. 8.**
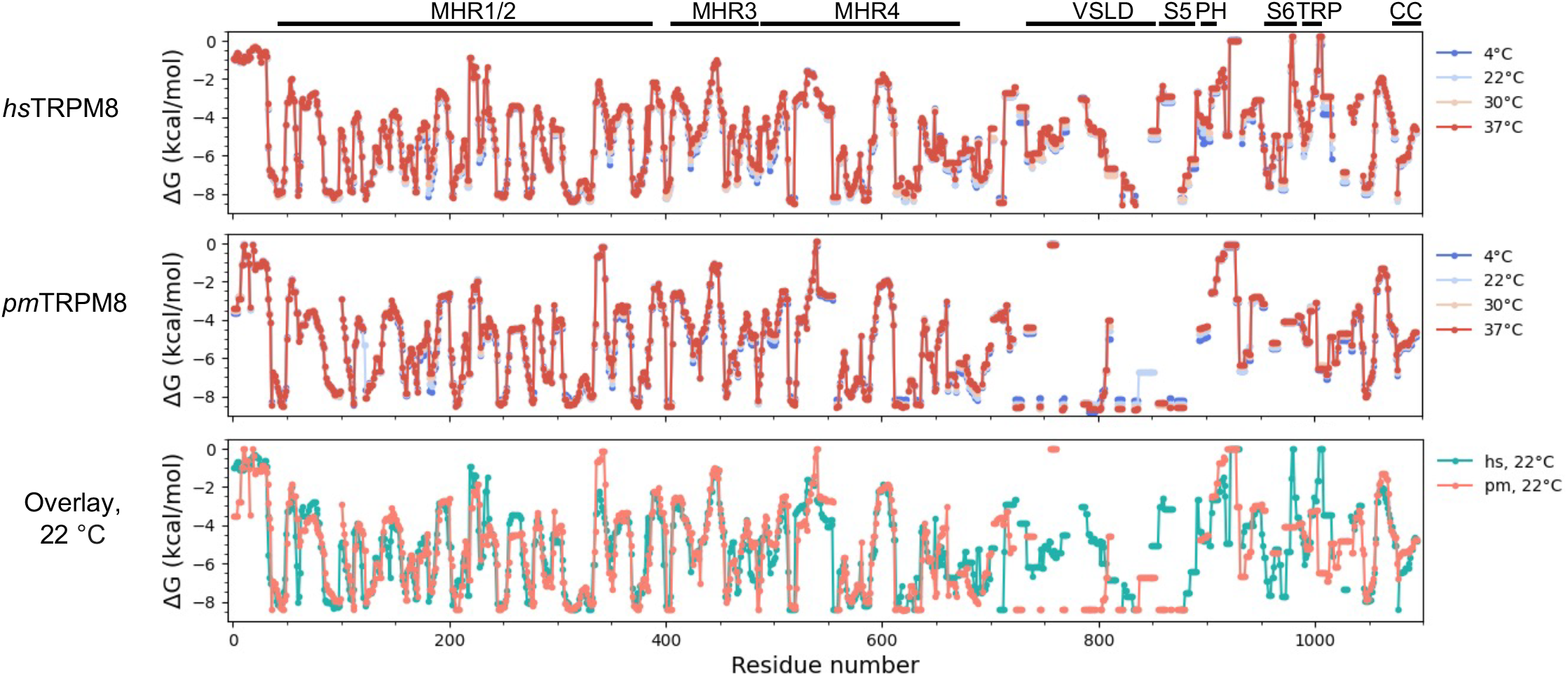
Comparison of the energetic profiles of human and avian TRPM8 at different temperatures. Residue numbers correspond to the avian sequence.

